# Functional Connectivity of Corticostriatal Circuitry and Psychosis-Like Experiences in the General Community

**DOI:** 10.1101/437376

**Authors:** Kristina Sabaroedin, Jeggan Tiego, Linden Parkes, Francesco Sforazzini, Amy Finlay, Beth Johnson, Ari Pinar, Vanessa Cropley, Ben J Harrison, Andrew Zalesky, Christos Pantelis, Mark Bellgrove, Alex Fornito

## Abstract

**Background:** Psychotic symptoms are proposed lie on a continuum, ranging from isolated psychosis-like experiences (PLEs) in non-clinical populations to frank disorder. Here, we investigate neurobiological correlates of this symptomatologic continuum by examining whether functional connectivity of dorsal corticostriatal circuitry, which is disrupted in patients and high-risk individuals, is associated with the severity of subclinical PLEs.

**Methods:** A community sample of 672 adults with no history of psychiatric or neurological illnesses completed a battery of seven questionnaires spanning various PLE domains. Principal component analysis (PCA) estimated major dimensions of PLEs from the questionnaires. PCA dimension scores were then correlated with whole-brain voxelwise functional connectivity (FC) maps of the striatum in a subset of 353 participants who completed a resting-state neuroimaging protocol.

**Results:** PCA identified two dimensions of PLEs accounting for 62.57% of variance in the measures, corresponding to positive and negative PLEs. Reduced FC between the dorsal striatum and prefrontal cortex correlated with higher positive PLEs. Negative PLEs correlated with increased FC between the dorsal striatum and visual and sensorimotor areas. In the ventral corticostriatal system, positive and negative PLEs were both associated with FC between the ventro-rostral putamen and sensorimotor cortices.

**Conclusions:** Consistent with past findings in patients and high-risk individuals, subthreshold positive symptomatology is associated with reduced FC of the dorsal circuit. These findings suggest that the connectivity of this circuit tracks the expression of psychotic phenomena across a broad spectrum of severity, extending from the subclinical domain to clinical diagnosis.

## Introduction

Psychotic symptoms are proposed to follow a continuous distribution of severity, ranging from the absence of symptoms at one end to frank disorder at the other (1). Subthreshold psychosis-like experiences (PLEs) lie between these extremes (2). Typically characterized as attenuated (i.e. subclinical or subthreshold) forms of canonical positive symptoms (e.g. delusions and hallucinations), PLEs also encompass subclinical variation of the negative symptoms of schizophrenia such as mild anhedonia (3–7). PLEs are predominantly transient (8, 9), and their persistent expression is thought to reflect an enduring personality dimension (10). The prevalence of PLEs in the general community reaches up to 8% (11), with higher incidences in the relatives of schizophrenia patients (12), suggesting that one’s liability for PLEs tracks vulnerability to clinical disorder. Severe PLEs may indeed progress into a help-seeking phase, as exemplified by the at-risk mental state (ARMS) for psychosis (13–17).

The continuum model of psychosis severity accords with evidence for a polygenic contribution to schizophrenia liability, which predicts a continuous population distribution of symptomatology (18, 19). It is also supported by neuroimaging evidence suggesting that subclinical and clinical phenomena share common neural correlates. The severity of PLEs in non-clinical samples correlates with variations in brain systems implicated in schizophrenia and various psychotic disorders, including frontotemporal, default mode, and cinguloopercular systems (20–30). Alterations of white matter integrity in the corpus callosum, thalamus, parietal, language and visual areas have also been commonly reported in studies of clinical and non-clinical individuals (31–33). These findings suggest that a continuum of neural function may underlie a broad spectrum of symptom expression.

The brain’s corticostriatal (CST) circuits are particularly relevant to the various cognitive and symptom dimensions of schizophrenia (34, 35). These parallel yet integrated circuits topographically connect frontal regions with the striatum, with feedback loops passing through the pallidum and thalamus (36, 37). The two most relevant CST circuits to psychosis are the ventral and dorsal systems (38, 39). The ventral, so-called limbic, system connects the orbitofrontal cortex, medial prefrontal cortex (PFC), and limbic structures of the brain (e.g. the hippocampus and amygdala) with the ventral striatum (36, 37, 40). This system is a major pathway for mesolimbic dopamine (41), and has long been implicated in psychosis due to its role in reward processing and associative learning (42).

The dorsal, so-called associative or cognitive system, links the dorsolateral prefrontal cortex (DLPFC) with the dorsal striatum (37). Positron emission tomography (PET) studies indicate that both dopamine synthesis capacity and synaptic concentration are elevated prominently in the dorsal striatum of schizophrenia patients (43), ARMS individuals (44, 45), and healthy people with increased liability for psychosis due to either genetic or environmental factors (46, 47). In ARMS, these elevations are only present in individuals who later transition to psychosis (48). These PET findings are complemented by studies of striatal functional connectivity, which report reduced coupling of the DLPFC with dorsal caudate and putamen in first-episode psychosis patients, their unaffected first-degree relatives, and ARMS individuals (49, 50). Similar changes have been found in first-episode mania patients with psychosis (51), suggesting that dorsal CST dysfunction tracks the emergence of psychotic symptoms across diagnostic categories. Other studies focusing on thalamic connectivity support the importance of dorsal CST function in risk for psychosis (52–54). Dorsal CST changes have been correlated with the severity of both positive and negative symptoms in ARMS and clinical groups (49, 50, 53), and are often accompanied by increased coupling in the ventral CST circuit and thalamic sensorimotor systems (49, 52–54).

Together, these results indicate that reduced functional coupling of the dorsal CST system tracks the severity of psychotic symptom expression across a wide spectrum of disease risk that spans genetic high-risk, ARMS, and clinically diagnosed individuals (38). Here, we investigated whether variation in CST function also tracks subclinical variation in PLEs in a large, non-clinical sample. We combined resting-state functional magnetic resonance imaging (fMRI) with an extensive battery of PLE questionnaires measuring a broad array of subclinical phenomena related to schizophrenia symptomatology. Following evidence in clinical and high-risk individuals (49, 50), we hypothesized that reduced coupling between the dorsal striatum and prefrontal cortex would be associated with more severe PLEs, particularly those related to the positive symptoms of schizophrenia. Secondarily, we evaluated whether increased coupling in the ventral system would correlate with more severe PLEs, as suggested by work in genetic high-risk individuals (49).

## Methods and Materials

### Participants

We recruited 672 participants (274 males; age range = 18-50 years old, mean [SD] age = 23.2 [4.89]) from the general community to complete an online battery of PLE measures. All participants were right handed with no personal history of neurological or psychiatric illness and no significant drug use (see Supplemental Information for further details). Recruitment was part of a larger genetics study that required participants to have all four grandparents of European descent. The study was conducted in accordance with the Monash University Human Research Ethics Committee (reference number 2012001562). Each participant provided written informed consent following a thorough explanation of the study.

A subset of 379 participants with complete PLE measures underwent our resting-state fMRI protocol. Participants were subsequently excluded for either scan artifacts, poor scan quality, or excessive in-scanner head motion (details in Supplemental Information). The final sample with complete PLE measures and neuroimaging data comprised 353 participants (155 males; median age [range] = 22 [18-50 years]; IQ range = 81-139, mean [SD] = 112 [11.6]).

### Measures of Psychosis-Like Experiences

To sample a wide range of variation in PLEs, we used seven psychometrically validated self-report measures of subthreshold psychotic symptoms: the short-form Oxford-Liverpool Inventory of Feelings and Experiences (sO-LIFE) (55), the Peter Delusion Inventory (PDI-21) (56), the Community Assessment of Psychotic Experience (CAPE) (5), and four Chapman Scales measuring magical ideation, perceptual aberration, and social and physical anhedonia (57–59). For the PDI and CAPE, subscales measuring distress and frequency were excluded from further analysis as they were redundant with the severity scales (all ρ >.9). The battery yielded a total of 272 items spanning 12 subscales (Supplemental Table S1).

### Principal Component Analysis

We used principal component analysis (PCA) of the subscale scores, with Varimax rotation, as implemented in IBM SPSS version 25, to derive data-driven estimates of the latent dimensions driving PLE variance in our sample. PCA was performed on the larger sample of 672 participants to obtain robust estimates of latent dimensions. The Kaiser-Meyer-Olkin test of sampling adequacy value was 0.87, indicating that the correlations between variables would yield reliable factors (60). Following PCA, component scores were extracted for all participants using the Anderson-Rubin method to ensure orthogonality (61).

### Neuroimaging Data Acquisition and Preprocessing

Multiband resting-state echo-planar images (EPI; 620 volumes, 754 milliseconds repetition time, 3mm isotropic voxels) and anatomical T1-weighted scans (1mm isotropic voxels) were acquired for each participant. The following pre-preprocessing steps were applied to the EPI images: (1) basic preprocessing in FSL FEAT which included removal of the first four volumes, rigid-body head motion correction, 3mm spatial smoothing, and high-pass temporal filter (75 seconds cut-off); (2) removal of artifacts using FSL-FIX; (3) spatial normalization to the MNI152 template; and (4) spatial smoothing with a 6mm full-width half-maximum Gaussian kernel. After processing, the data were subjected to rigorous quality control for motion artifacts, as per past work (62). Further details are in Supplemental Information.

### Definition of Seed Regions of Interest

In each hemisphere, six striatal regions-of-interest (ROI) were seeded using a 3.5mm radius spheres, as per past work (49, 50, 63). For the caudate, three ROIs were seeded along a dorsoventral axis, including the dorsal caudate (DC; x = ±13, y = 15, z = 9), the superior ventral caudate (x = ±10, y = 15, z = 0), and the inferior ventral caudate/nucleus accumbens (x = ±9, y = 9, z = −8). Three ROIs were seeded for the putamen along a similar axis, comprising the dorso-caudal putamen (DCP; x = ±28, y = 1, z = 3), the dorso-rostral putamen (DRP; x = ±25, y = 8, z = 6), and the ventro-rostral putamen (VRP; x = ±20, y = 12, z = −3). Seeds in the dorsal CST system comprise the DC, DRP, and DCP; whereas seeds in the ventral system comprise the inferior ventral caudate/nucleus accumbens, superior ventral caudate, and VRP. The mean time series of each region was then used for seed-related functional connectivity mapping.

### Functional Connectivity Analysis

First-level analysis was performed using SPM8 as previously described (49, 50). For each participant, a general linear model containing the six seed region time courses as covariates was used to model BOLD signal fluctuations in each voxel. Separate models were fitted for the left and right hemispheres, yielding a pair of pair of brain maps for each striatal ROI.

Parameter estimates from the first-level analysis were passed to a second-level general linear model to generate group-wide functional connectivity maps for each ROI. Covariates comprised component scores of orthogonal PLE dimensions derived from the PCA. Nuisance covariates included age, age^2^, IQ, sex, and mean framewise displacement as a measure of in-scanner motion (64). Correlations between PLEs and seed-related functional connectivity, collapsed across hemispheres, were declared significant if they passed a threshold-free cluster enhancement (TFCE) (65) correction of *p* < .05, determined using 5000 permutations, as implemented in FSL Randomise (66). Scatterplots of associations between PLEs and functional connectivity were visualized using a leave-one-subject-out (LOSO) approach that circumvents circular inference (67) (details in Supplemental Information).

## Results

### Principal Component Analysis

PCA was performed on 12 subscales derived from seven PLE questionnaires completed by 672 participants (descriptive statistics and pairwise correlations between subscales are in Supplement Table S1). Based on the inflection in the scree plot (Supplement Figure S1) (68), we retained two principal components. The first component accounted for 42.87% of the variance, with high loadings from subscales measuring the positive dimension of psychosis-related experiences including delusional ideation, unusual experiences, perceptual aberrations, and eccentric behavior (i.e. positive PLEs). The second component accounted for 19.7% of the variance, with high loadings from subscales measuring the negative dimension of psychosis-related experiences (i.e. negative PLEs), including social and physical anhedonia. Component loadings are displayed in Table 1.

**Table 1.**
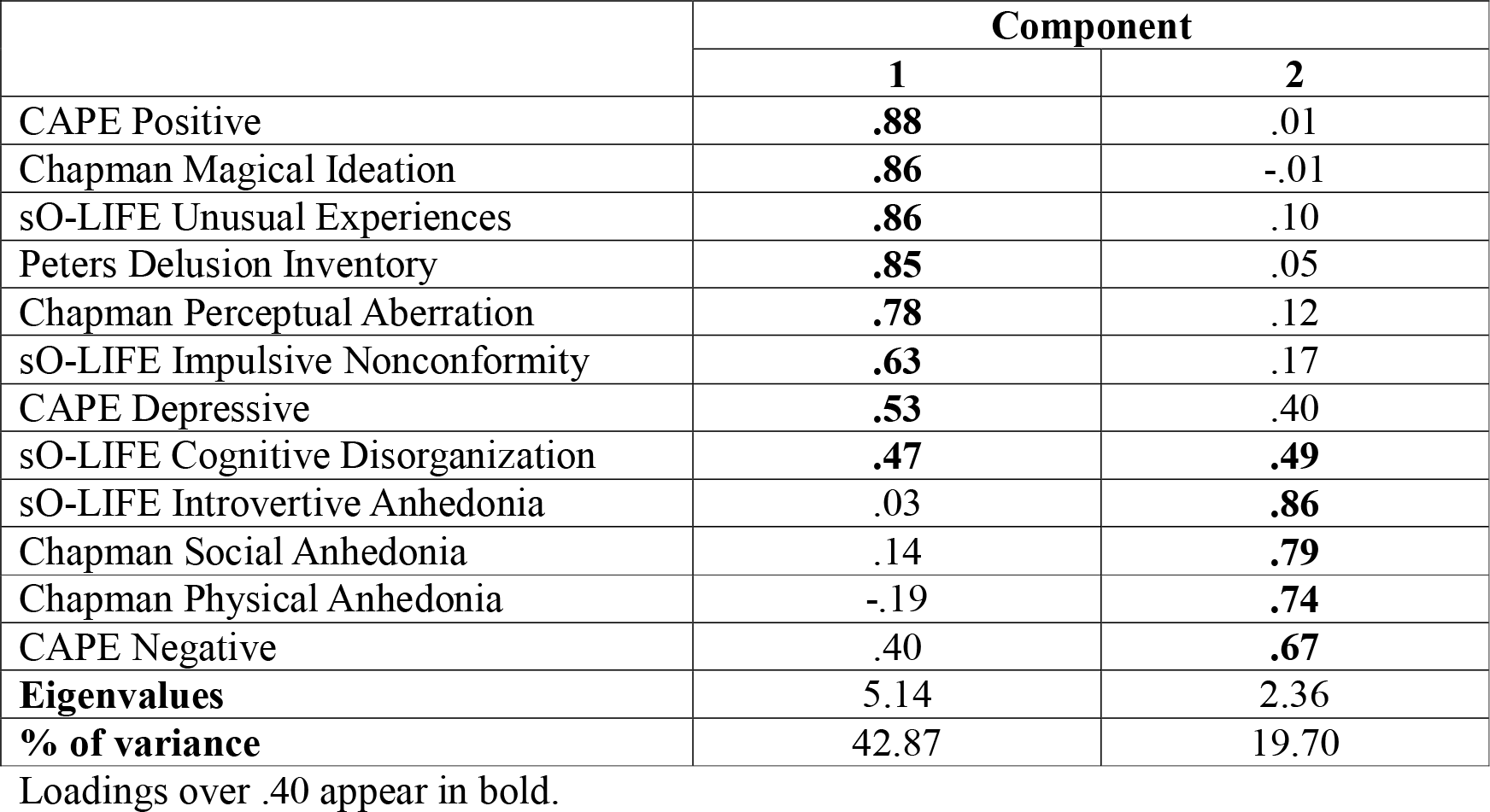
Component Loadings of Subscales After Rotation.

### Corticostriatal Functional Connectivity

Functional connectivity analysis was performed in 353 participants with fMRI data. Each striatal region showed functional connectivity profiles consistent with known anatomy and prior findings (Supplement Figure S3) (49, 50, 63).

#### Dorsal CST functional connectivity and PLEs

As predicted, higher scores on the positive PLE dimension were associated with reduced functional connectivity between the DRP and the right DLPFC (i.e. anterior middle frontal gyrus) (Table 2; Figure 1). Higher positive PLE scores were also associated with reduced coupling between the DC and left dorsal anterior cingulate cortex (ACC); reduced coupling between the DCP and left visual cortex, left posterior cingulate sulcus, and the right primary motor and sensory regions (Table 2; Figure 1). Higher positive PLE scores were also linked with increased coupling between the DCP and regions of right medial PFC, frontal eye fields and frontal pole; and increased coupling between the DC and the lateral occipital cortex (Table 2).

**Table 2.**
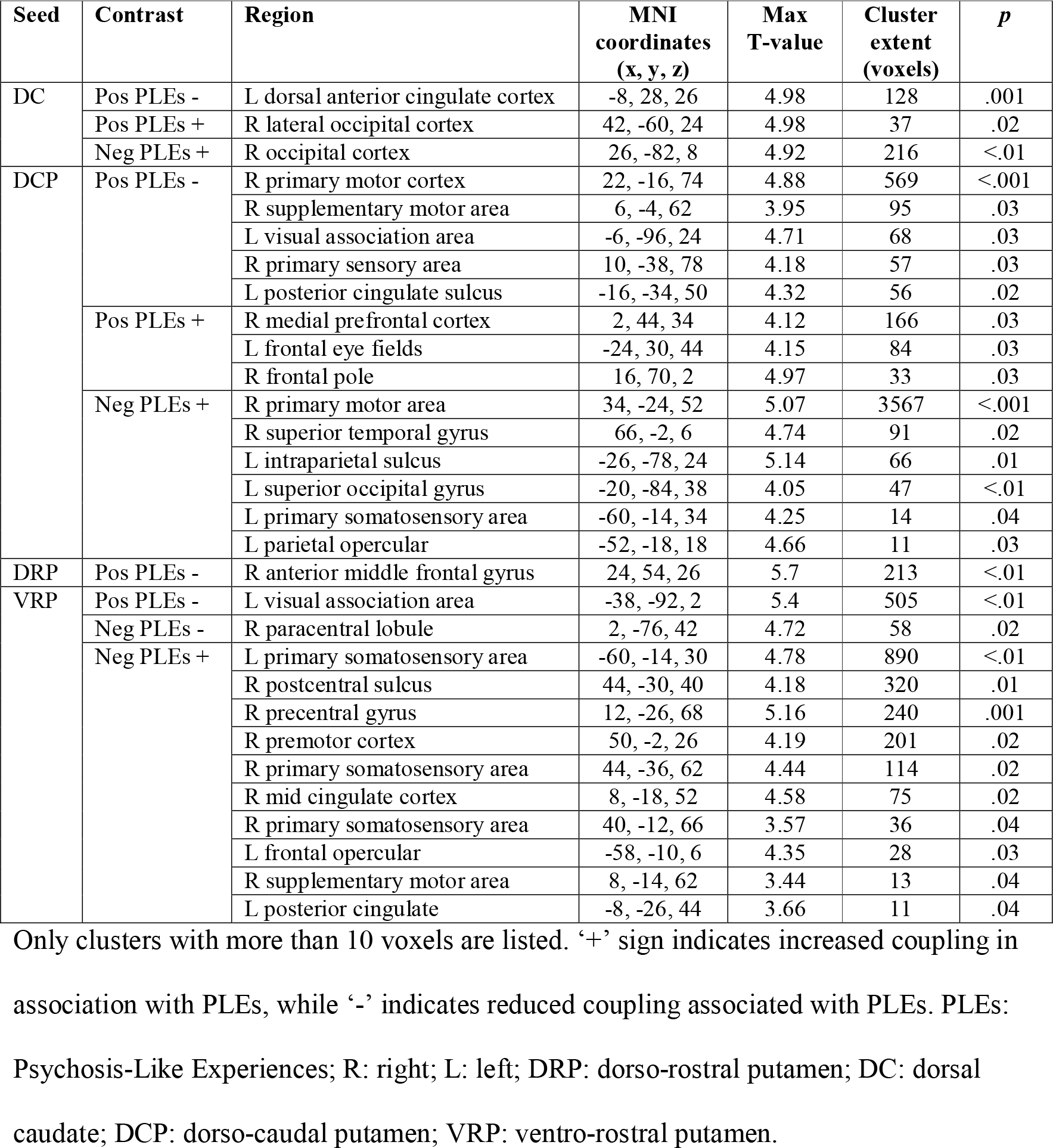
Regions Where Striatal Functional Connectivity Was Associated with Psychosis-Like Experiences (*p* < .05 TFCE Corrected)

**Figure 1.**
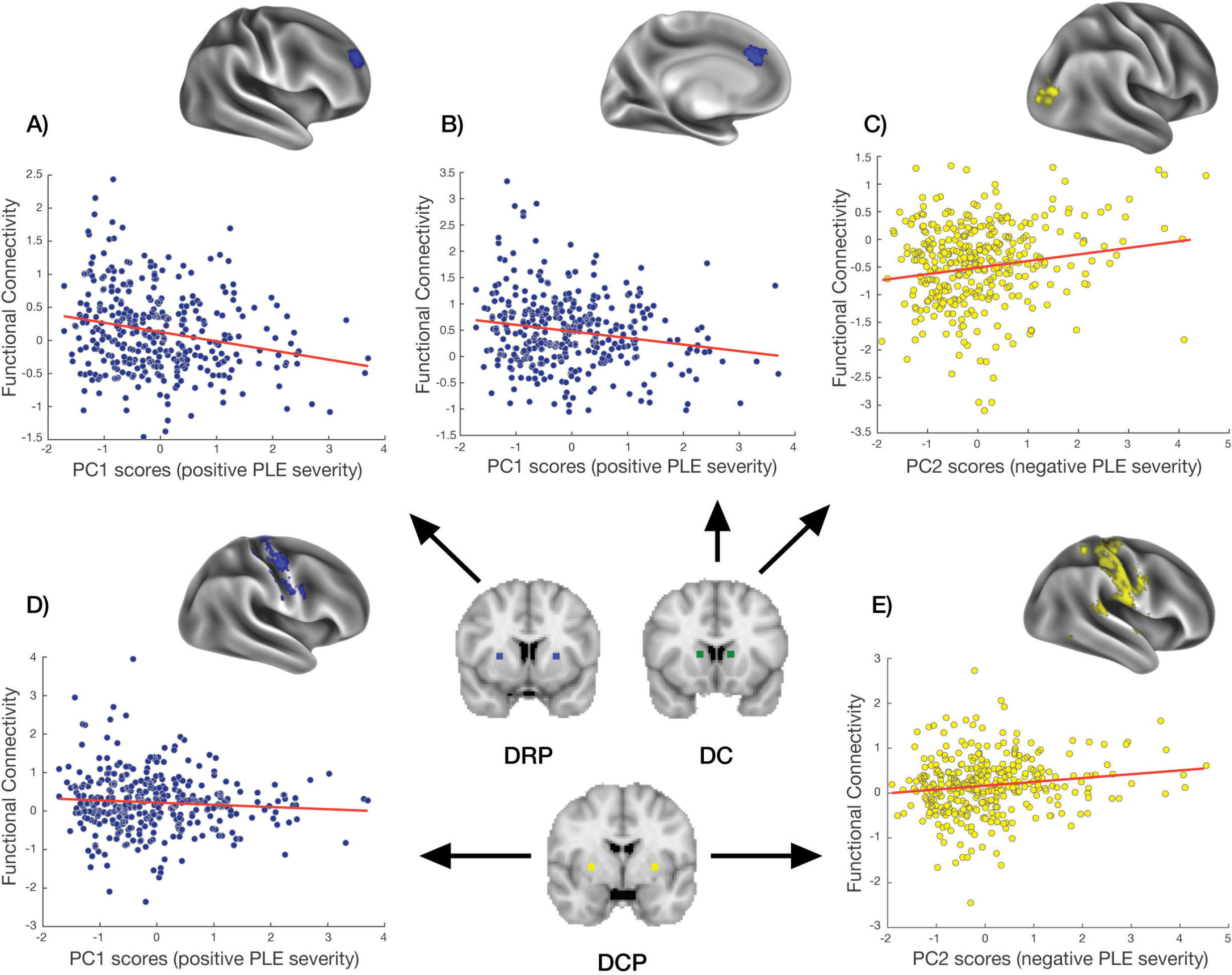
Associations Between Dorsal Circuit Functional Connectivity and Psychosis-Like Experiences. Coronal slices in the center of the bottom row depict the location of striatal seed regions in the dorsal circuit. Cortical surface maps depict the cortical sites for which functional connectivity with each seed correlates with PLE severity. Blue scatterplots depict the associations between positive PLE severity and (A) DRP and right DLPFC functional coupling (B) DC and left mPFC coupling; and (D) DCP and right sensorimotor coupling. Yellow scatterplots depict the associations between negative PLE severity and (C) DC and right visual cortex coupling and (E) DCP and right sensory cortex coupling. Clusters are thresholded at *p* < .05 TFCE corrected. For visualization purposes, functional connectivity estimates were obtained using Leave-One-Subject-Out analysis. DRP: dorso-rostral putamen; DC: dorsal caudate; DCP: dorso-caudal putamen; PC: principal component; PLEs: Psychosis-Like Experiences.

Higher scores on the negative PLE dimension were associated with increased functional connectivity between the dorsal striatum and sensorimotor areas; namely, between the DCP and right primary motor cortex, right superior temporal gyrus, and left occipital and somatosensory areas; and between the DC seed and right occipital cortex (Table 2; Figure 1).

#### Ventral CST functional connectivity and PLEs

Associations between PLEs and ventral system functional connectivity were limited to the VRP seeds. Specifically, higher positive PLEs were associated with lower functional connectivity between this region and left visual cortex (Table 2; Figure 2). Higher negative PLEs were associated with higher coupling between the VRP and bilateral sensorimotor cortex (Table 2; Figure 2), and lower coupling between the VRP and the right paracentral lobule (Table 2).

**Figure 2.**
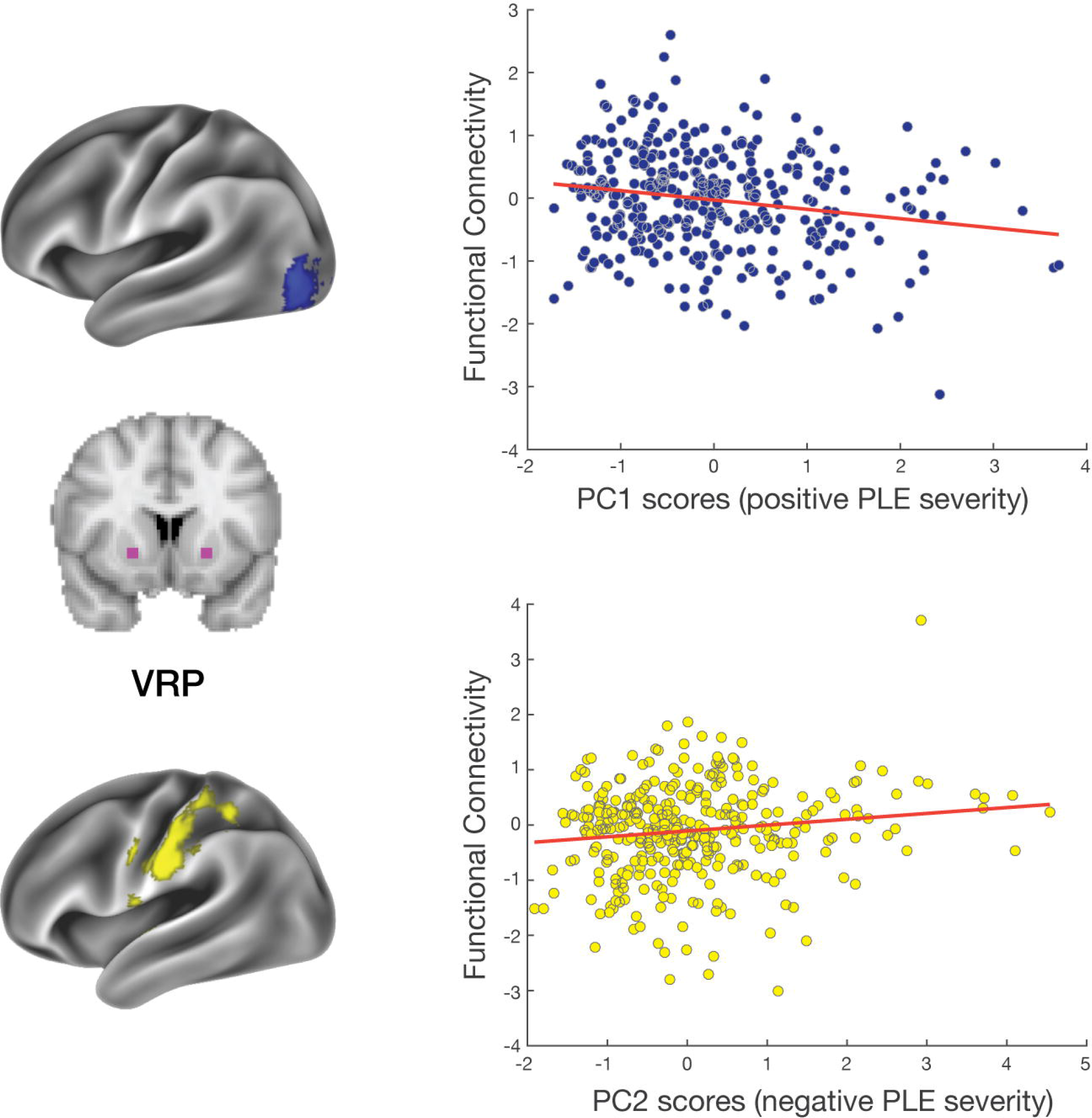
Associations Between Ventral Circuit Functional Connectivity and Psychosis-Like Experiences. Coronal slice in the center of the left column depicts the location of striatal seed regions in the ventral circuit. Cortical surface maps show the cortical sites for which functional connectivity with the ventral seeds correlate with PLE severity. Blue scatterplot depicts the association between positive PLE and the functional coupling of VRP and left visual area. Yellow scatterplot depicts the association between negative PLEs and VRP and left somatosensory area. Clusters are thresholded as *p* < .05 TFCE corrected. For visualization purposes, functional connectivity estimates were obtained using a Leave-One-Subject-Out analysis. VRP: ventro-rostral putamen; PC: principal component; PLEs: Psychosis-Like Experiences.

## Discussion

Corticostriatal systems have long been implicated in the pathophysiology of psychosis. Converging evidence from studies in independent samples report that reduced coupling of the dorsal CST circuit is apparent across a broad spectrum of disease risk (49, 50, 52–54). Here, we show that reduced functional coupling in the dorsal CST circuit tracks subclinical variation in PLEs related to positive symptomatology, consistent with a continuum of neural function that tracks the severity of symptom expression (2, 10, 11) (see Figure 3 for a summary). Our comprehensive investigation of striatal functional connectivity identified several other associations between PLEs and CST coupling that have not been observed in patients, suggesting a discontinuity for these phenotypes.

**Figure 3.**
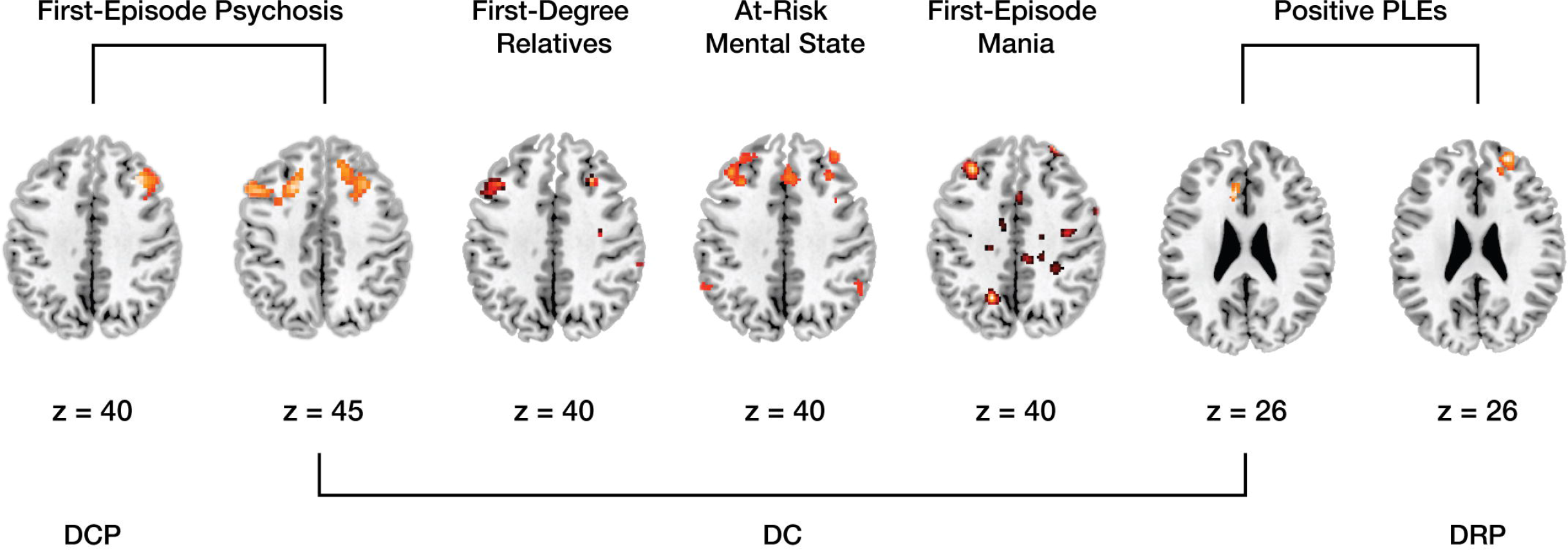
Functional Connectivity of the Dorsal Circuit Across a Broad Spectrum of Disease Risk. Axial slices showing prefrontal regions where coupling within the dorsal circuit was reduced across a broad spectrum of disease risk, as identified in the current study and past findings. From left to right: regions in which coupling with the DCP and DC were reduced in first-episode psychotic patients compared to healthy controls (data from reference (49)); regions of reduced coupling with the DC in healthy first-degree relatives of patients, ARMS individuals and first-episode mania patients with psychosis compared to healthy controls (data from references (49–51)); and findings from the present study, showing regions where lower coupling with the DC and DRP was associated with more sever positive PLEs. The z-axis slice in MNI coordinates is shown beneath each image. Left hemisphere is on the left. DCP: dorso-caudal putamen; DC: dorsal caudate; DRP: dorso-rostral putamen.

### Dorsal Corticostriatal Coupling and PLEs

As hypothesized, higher levels of positive PLEs were associated with reduced coupling between the dorsal striatum and PFC. This association is broadly consistent with our past work in patients, genetic high-risk, and ARMS individuals (49, 50). However, the specific regions implicated in this past work show some differences with our current findings. Specifically, we have previously reported that first-episode patients, their unaffected relatives, and ARMS individuals show reduced coupling between the dorsal caudate and DLPFC, with patients and relatives also showing reduced coupling between the dorso-caudal putamen and DLPFC (49, 50). Here, we found that positive PLEs were associated with reduced coupling between the dorsal caudate and ACC, and between dorso-*rostral* putamen and a more anterior region of DLPFC (Figure 3).

One explanation for these discrepancies is that variations in sample characteristics and image processing techniques may lead to slight changes in the localization of clinically relevant effects. Another possibility is that our findings reflect a weak continuum model, in which PLEs are broadly related to the activity of the dorsal CST system, but the onset of frank illness only arises with dysfunction in very specific elements of this system (putatively involving the dorsal caudate and DLPFC). It is also possible that our use of PCA to measure PLE severity, while capturing the dominant modes of variance across a wide battery of measures, may miss a more specific component of positive symptomatology that shows high behavioral and neurobiological continuity with clinical disorder. Better models of the latent dimensions underlying psychosis-related psychopathology across the full range of illness severity will be required to clarify the relationship between clinical and subclinical phenotypes.

Other associations with dorsal circuit function and PLE severity implicate areas that are known to be affected by schizophrenia and subthreshold symptomatology. Abnormal structure and function of the ACC are frequently reported in both patients and high-risk individuals (29, 69–74). Concordantly, ACC dysfunction is linked to a failure to update prior beliefs based on sensory experience, which may induce hallucinations (75). Positive PLE severity was also associated with increased coupling between the dorsal striatum and regions involved in visual processing and oculomotor control which are processes that are disrupted in schizophrenia (36, 76, 77). Our findings of reduced coupling between the dorsal seeds and the PFC in association with higher positive PLEs are consistent with a report of simultaneously decreased and increased activation in the dorsal striatum and PFC, respectively, in healthy adolescents with high PLEs during a reward processing task (78).

Negative PLE severity was generally associated with increased dorsal circuit coupling. The implicated regions involve the dorsal caudate, dorso-caudal putamen, and sensorimotor systems. This finding is distinct from our prior report that negative symptoms correlated with reduced coupling between the dorsal caudate and DLPFC in first-episode patients (49). Negative symptoms are complex and can be difficult to assess via self-report, with such measures varying in their sensitivity to quantifying different constructs of the dimension (79). Detailed investigation of the concordance between self-report and clinician-rated assessments of negative symptoms in both clinical and subclinical domains will be required to further refine this phenotype.

### Ventral Corticostriatal Coupling and PLEs

Associations between PLEs and ventral CST coupling were restricted to the VRP-sensorimotor and visual circuit, such that increased functional connectivity was associated with higher negative and lower positive PLE scores. The association with positive PLEs is consistent with disrupted fronto-striatal connectivity leading to increased thalamic outflow to sensory cortices (38, 80). The concomitant association with negative PLEs implies that this increased outflow may be accompanied by psychomotor difficulties, which characterize many negative symptoms of schizophrenia (81–83), given that CST circuits modulate goal directed behavior involving immediate physical action by coordinating motor with executive functions (83).

In contrast with our prior report that first-episode psychosis patients and their unaffected relatives showed increased coupling between the nucleus accumbens and ventral frontal cortex (49), we found no evidence for an association between the connectivity of this system and PLE severity. There is some evidence to suggest that this system may be tied to specific aspects of negative PLEs such as social anhedonia in subclinical groups (84), which may not have been identified in our analysis that focused on dominant modes of PLE variance.

### Limitations

Over 90% of our sample was aged under 30 years, hence most participants have not passed through the maximal period of risk for schizophrenia (85). As psychotic experiences in early adulthood have been found to predict later psychopathology (15, 86), it is possible that some people in our cohort may develop the illness at a later stage. Our exclusion of individuals with a personal history of mental health treatment, ensured that we sampled the subclinical range of symptom expression, but this means that we may not have sampled the more severe end of the PLE spectrum.

Our two-component PCA solution parallels the well-known classification of clinical positive and negative symptomatology, but should be interpreted with some caveats. First, as stated, the leading components from the PCA encompass common variance across measures and may miss subtle aspects of PLEs that are unique to each questionnaire. Second, the simple two-factor solution may reflect an implicit bias in the PLE questionnaires to (over-)sample experiences related to positive and negative symptom dimensions of psychotic illness, which may inadvertently exclude other dimensions. A comprehensive assessment of psychotic and non-psychotic symptomatology in subclinical populations may lead to a more refined model of the psychosis risk phenotype.

Our scatterplots show small-to-moderate associations between PLEs and coupling estimates. We note that these were generated using a LOSO analysis for visualization purposes to avoid circular inference (67), and do not directly correspond to the general linear model used to estimate the main functional connectivity results. Finally, in-scanner head motion exerts a pernicious effect on functional connectivity estimates (62, 64, 87, 88), but our extensive quality control procedures indicate that our findings could not be explained by motion artifact (see Supplemental Information).

### Conclusions

Our findings indicated that variation in dorsal CST function tracks subclinical expression of PLEs in a non-clinical sample, paralleling the circuit-level changes seen in patients and high-risk groups. Together, these findings are consistent with a continuum of psychosis risk that is apparent at the level of overt behavior and underlying neurobiology, and which spans across a broad spectrum of liability ranging from isolated experiences or attributional biases in otherwise healthy individuals, to frank disorder in clinical populations. Early interventions targeting the function of this circuit may thus be a fruitful strategy for mitigating disease risk.

## Author Affiliations

Brain and Mental Health Research Hub, Monash Institute of Cognitive and Clinical Neurosciences, Monash University, Victoria, Australia (Sabaroedin, Tiego, Parkes, Finlay, Fornito); School of Psychological Sciences, Monash Institute of Cognitive and Clinical Neurosciences, Monash University, Victoria, Australia (Sabaroedin, Tiego, Parkes, Finlay, Johnson, Pinar, Bellgrove, Fornito); Monash Biomedical Imaging, Monash University, Victoria, Australia (Sforazzini, Fornito); Melbourne Neuropsychiatry Centre, Department of Psychiatry, The University of Melbourne, Victoria, Australia (Cropley, Harrison, Zalesky, Pantelis)

## Author Contributions

Sabaroedin and Fornito had full access to all the data in the study and takes responsibility for the integrity of the data and the accuracy of data analysis

*Study concept and design:* Fornito, Bellgrove, Harrison, Zalesky, Pantelis

*Drafting of manuscript:* Sabaroedin, Fornito

*Acquisition of Data:* Tiego, Finlay, Johnson, Pinar, Sabaroedin, Bellgrove, Fornito

*Analysis and interpretation of data:* Sabaroedin, Fornito, Tiego, Parkes, Sforazzini

*Critical revision of manuscript for important intellectual content:* Bellgrove, Tiego, Cropley, Harrison, Zalesky, Pantelis

*Statistical analysis:* Sabaroedin, Tiego, Fornito

*Administrative, technical, or material support:* Tiego, Finlay, Johnson

*Supervision:* Bellgrove, Fornito

## References

1. Grant P, Green MJ, Mason OJ (2018): Models of schizotypy: the importance of conceptual clarity. Schizophr Bull. 1–8.

2. Kelleher I, Cannon M (2011): Psychotic-like experiences in the general population: characterizing a high-risk group for psychosis. Psychol Med. 41: 1–6.

3. Barrantes-Vidal N, Chun CA, Myin-Germeys I, Kwapil TR (2013): Psychometric schizotypy predicts psychotic-like, paranoid, and negative symptoms in daily life. J Abnorm Psychol. 122: 1077–1087.

4. Verdoux H, Van Os J (2002): Psychotic symptoms in non-clinical populations and the continuum of psychosis. Schizophr Res. 54: 59–65.

5. Stefanis NC, Hanssen M, Smirnis NK, Avramopoulos DA, Evdokimidis IK, Stefanis CN, et al. (2002): Evidence that three dimensions of psychosis have a distribution in the general population. Psychol Med. 32: 347–358.

6. MacDonald AW, Pogue-Geile MF, Debski TT, Manuck S (2001): Genetic and environmental influences on schizotypy: a community-based twin study. Schizophr Bull. 27: 47–58.

7. Linney YM, Murray RM, Peters ER, MacDonald AM, Rijsdijk F, Sham PC (2003): A quantitative genetic analysis of schizotypal personality traits. Psychol Med. 33: 803–816.

8. Hanssen M, Krabbendam L, Vollema M, Delespaul P, van Os J (2006): Evidence for instrument and family-specific variation of subclinical psychosis dimensions in the general population. J Abnorm Psychol. 115: 5–14.

9. Cougnard A, Marcelis M, Myin-Germeys I, De Graaf R, Vollebergh W, Krabbendam L, et al. (2007): Does normal developmental expression of psychosis combine with environmental risk to cause persistence of psychosis? A psychosis proneness-persistence model. Psychol Med. 37: 513–527.

10. Ettinger U, Meyhöfer I, Steffens M, Wagner M, Koutsouleris N (2014): Genetics, cognition, and neurobiology of schizotypal personality: a review of the overlap with schizophrenia. Front Psychiatry. 5: 1–16.

11. van Os J, Linscott RJ, Myin-Germeys I, Delespaul P, Krabbendam L (2009): A systematic review and meta-analysis of the psychosis continuum: evidence for a psychosis proneness-persistence-impairment model of psychotic disorder. Psychol Med. 39: 179–95.

12. Vollema MG, Sitskoorn MM, Appels MC, Kahn RS (2002): Does the Schizotypal Personality Questionnaire reflect the biological-genetic vulnerability to schizophrenia? Schizophr Res. 54: 39–45.

13. Cannon TD, Cornblatt B, McGorry P (2007): Editor’s introduction: The empirical status of the ultra high-risk (prodromal) research paradigm. Schizophr Bull. 33: 661–664.

14. Kaymaz N, Drukker M, Lieb R, Wittchen HU, Werbeloff N, Weiser M, et al. (2012): Do subthreshold psychotic experiences predict clinical outcomes in unselected non-help-seeking population-based samples? A systematic review and meta-analysis, enriched with new results. Psychol Med. 42: 2239–2253.

15. Werbeloff N, Drukker M, Dohrenwend BP, Levav I, Yoffe R, van Os J, et al. (2012): Self-reported attenuated psychotic symptoms as forerunners of severe mental disorders later in life. Arch Gen Psychiatry. 69: 467–475.

16. Poulton R, Caspi A, Moffitt TE, Cannon M, Murray RM, Harrington H (2000): Children’s self-reported psychotic symptoms and adult schizophreniform disorder: a 15-year longitudinal study. Arch Gen Psychiatry. 57: 1053–1058.

17. Yung AR, McGorry PD (1996): The prodromal phase of first-episode psychosis: past and current conceptualizations. Schizophr Bull. 22: 353–370.

18. Lee SH, DeCandia TR, Ripke S, Yang J, Sullivan PF, Goddard ME, et al. (2012): Estimating the proportion of variation in susceptibility to schizophrenia captured by common SNPs. Nat Genet. 44: 247–50.

19. Plomin R, Haworth CM, Davis OSP (2009): Quantitative Traits. Nat Rev Genet. 10: 872–878.

20. Ettinger U, Williams SCR, Meisenzahl EM, Möller HJ, Kumari V, Koutsouleris N (2012): Association between brain structure and psychometric schizotypy in healthy individuals. World J Biol Psychiatry. 13: 544–549.

21. Satterthwaite TD, Vandekar SN, Wolf DH, Bassett DS, Ruparel K, Shehzad Z, et al. (2015): Connectome-wide network analysis of youth with Psychosis-Spectrum symptoms. Mol Psychiatry. 20: 1508–1515.

22. Henseler I, Falkai P, Gruber O (2010): Disturbed functional connectivity within brain networks subserving domain-specific subcomponents of working memory in schizophrenia: relation to performance and clinical symptoms. J Psychiatr Res. 44: 364–372.

23. Sheffield JM, Kandala S, Burgess GC, Harms MP, Barch DM (2016): Cingulo-opercular network efficiency mediates the association between psychotic-like experiences and cognitive ability in the general population. Biol Psychiatry Cogn Neurosci Neuroimaging. 1: 498–506.

24. Wolthusen RPF, Coombs G, Boeke EA, Ehrlich S, DeCross SN, Nasr S, Holt DJ (2018): Correlation between levels of delusional beliefs and perfusion of the hippocampus and an associated network in a non-help-seeking population. Biol Psychiatry Cogn Neurosci Neuroimaging. 3: 178–186.

25. Fusar-Poli P, Broome MR, Matthiasson P, Woolley JB, Johns LC, Tabraham P, et al. (2010): Spatial working memory in individuals at high risk for psychosis: longitudinal fMRI study. Schizophr Res. 123: 45–52.

26. Corlett PR, Fletcher PC (2012): The neurobiology of schizotypy: fronto-striatal prediction error signal correlates with delusion-like beliefs in healthy people. Neuropsychologia. 50: 3612–3620.

27. Garrity AG, Pearlson GD, McKiernan K, Lloyd D, Kiehl KA, Calhoun VD (2007): Aberrant “default mode” functional connectivity in schizophrenia. Am J Psychiatry. 164: 450–457.

28. Whitfield-Gabrieli S, Thermenos HW, Milanovic S, Tsuang MT, Faraone S V., McCarley RW, et al. (2009): Hyperactivity and hyperconnectivity of the default network in schizophrenia and in first-degree relatives of persons with schizophrenia. Proc Natl Acad Sci. 106: 1279–1284.

29. White TP, Joseph V, Francis ST, Liddle PF (2010): Aberrant salience network (bilateral insula and anterior cingulate cortex) connectivity during information processing in schizophrenia. Schizophr Res. 123: 105–115.

30. Talati P, Rane S, Kose S, Blackford JU, Gore J, Donahue MJ, Heckers S (2014): Increased hippocampal CA1 cerebral blood volume in schizophrenia. NeuroImage Clin. 5: 359–364.

31. Jacobson S, Kelleher I, Harley M, Murtagh A, Clarke M, Blanchard M, et al. (2010): Structural and functional brain correlates of subclinical psychotic symptoms in 11-13 year old schoolchildren. Neuroimage. 49: 1875–1885.

32. Van Dellen E, Bohlken MM, Draaisma L, Tewarie PK, Van Lutterveld R, Mandl R, et al. (2016): Structural brain network disturbances in the psychosis spectrum. Schizophr Bull. 42: 782–789.

33. Skudlarski P, Schretlen DJ, Thaker GK, Stevens MC, Keshavan MS, Sweeney J a, et al. (2013): Diffusion tensor imaging white matter endophenotypes in patients with schizophrenia or psychotic bipolar disorder and their relatives. Am J Psychiatry. 170: 886–898.

34. Pantelis C, Barnes TRE, Nelson HE (1992): Is the concept of frontal-subcortical dementia relevant to schizophrenia? Br J Psychiatry. 160: 442–460.

35. Pantelis C, Brewer W (1996): Neurocognitive and neurobehavioural patterns and the syndromes of schizophrenia: role of frontal-subcortical networks. In: Pantelis C, Barnes T, He N, editors. Schizophr A Neuropsychol Perspect. London: John Wiley & Sons, pp 317–343.

36. Haber SN (2016): Corticostriatal circuitry. Dialogues Clin Neurosci. 18: 7–21.

37. Alexander G (1986): Parallel Organization of Functionally Segregated Circuits Linking Basal Ganglia and Cortex. Annu Rev Neurosci. 9: 357–381.

38. Dandash O, Pantelis C, Fornito A (2017): Dopamine, fronto-striato-thalamic circuits and risk for psychosis. Schizophr Res. 180: 48–57.

39. Grace AA (2016): Dysregulation of the dopamine system in the pathophysiology of schizophrenia and depression. Nat Rev Neurosci. 17.

40. Draganski B, Kherif F, Klöppel S, Cook PA, Alexander DC, Parker GJM, et al. (2008): Evidence for segregated and integrative connectivity patterns in the human basal ganglia. J Neurosci. 28: 7143–7152.

41. Haber SN, Knutson B (2009): The reward circuit: linking primate anatomy and human imaging. Neuropsychopharmacology. 35: 1–23.

42. Kapur S (2003): Psychosis as a state of aberrant salience: a framework linking biology, phenomenology, and pharmacology in schizophrenia. Am J Psychiatry. 160: 13–23.

43. Kegeles LS, Abi-Dargham A, Frankle WG, Gil R, Cooper TB, Slifstein M, et al. (2010): Increased synaptic dopamine function in associative regions of the striatum in schizophrenia. Arch Gen Psychiatry. 67: 231–239.

44. Howes OD, Montgomery AJ, Asselin MC, Murray RM, Valli I, Tabraham P, et al. (2009): Elevated striatal dopamine function linked to prodromal signs of schizophenia. Arch Gen Psychiatry. 66: 13–20.

45. Fusar-Poli P, Howes OD, Allen P, Broome M, Valli I, Asselin M (2010): Abnormal frontostriatal interactions in people with prodromal signs of psychosis: a multimodal imaging study. Arch Gen Psychiatry. 67: 683–691.

46. Egerton A, Howes OD, Houle S, McKenzie K, Valmaggia LR, Bagby MR, et al. (2017): Elevated striatal dopamine function in immigrants and their children: a risk mechanism for psychosis. Schizophr Bull. 43: 293–301.

47. Huttunen J, Heinimaa M, Svirskis T, Nyman M, Kajander J, Forsback S, et al. (2008): Striatal dopamine synthesis in first-degree relatives of patients with schizophrenia. Biol Psychiatry. 63: 114–117.

48. Howes O, Bose S, Turkheimer F, Valli I, Egerton a, Stahl D, et al. (2011): Progressive increase in striatal dopamine synthesis capacity as patients develop psychosis: a PET study. Mol Psychiatry. 16: 885–886.

49. Fornito A, Harrison BJ, Goodby E, Dean A, Ooi C, Nathan PJ, et al. (2013): Functional dysconnectivity of corticostriatal circuitry as a risk phenotype for psychosis. JAMA psychiatry. 70: 1143–51.

50. Dandash O, Fornito A, Lee J, Keefe RSE, Chee MWL, Adcock RA, et al. (2014): Altered striatal functional connectivity in subjects with an at-risk mental state for psychosis. Schizophr Bull. 40: 904–913.

51. Dandash O, Yücel M, Daglas R, Pantelis C, McGorry P, Berk M, Fornito A (2018): Differential effect of quetiapine and lithium on functional connectivity of the striatum in first episode mania. Transl Psychiatry. 8. doi: 10.1038/s41398-018-0108-8.

52. Anticevic A, Cole MW, Repovs G, Murray JD, Brumbaugh MS, Winkler AM, et al. (2014): Characterizing thalamo-cortical disturbances in schizophrenia and bipolar illness. Cereb Cortex. 24: 3116–3130.

53. Anticevic A, Haut K, Murray JD, Repovs G, Yang GJ, Diehl C, et al. (2015): Association of thalamic dysconnectivity and conversion to psychosis in youth and young adults at elevated clinical risk. JAMA psychiatry. 72: 882–891.

54. Woodward ND, Heckers S (2016): Mapping thalamocortical functional connectivity in chronic and early stages of psychotic disorders. Biol Psychiatry. 79: 1016–1025.

55. Mason O, Linney Y, Claridge G (2005): Short scales for measuring schizotypy. Schizophr Res. 78: 293–296.

56. Peters E, Joseph S, Day S, Garety P (2004): Measuring delusional ideation: the 21-Item Peters et al. Delusions Inventory. Schizophr Bull. 30: 1005–1022.

57. Eckblad M, Chapman LJ (1983): Magical ideation as an indicator of schizotypy. J Consult Clin Psychol. 51: 215–225.

58. Chapman LJ, Chapman JP, Raulin ML (1978): Body image aberration in schizophrenia. J Abnorm Psychol. 87: 399–407.

59. Chapman LJ, Chapman JP, Raulin ML (1976): Scales for physical and social anhedonia. J Abnorm Psychol. 85: 374–382.

60. Kaiser HF (1974): An index of factorial simplicity. Psychometrika. 39: 31–36.

61. DiStefano C, Zhu M, Mindrila D (2009): Understanding and using factor scores: considerations for the applied researcher. Pract Assessment, Res Eval. 14: 1–11.

62. Parkes L, Fulcher B, Yücel M, Fornito A (2018): An evaluation of the efficacy, reliability, and sensitivity of motion correction strategies for resting-state functional MRI. Neuroimage. 171: 415–436.

63. Di Martino A, Scheres A, Margulies DS, Kelly AMC, Uddin LQ, Shehzad Z, et al. (2008): Functional connectivity of human striatum: A resting state fMRI study. Cereb Cortex. 18: 2735–2747.

64. Power JD, Barnes KA, Snyder AZ, Schlaggar BL, Petersen SE (2012): Spurious but systematic correlations in functional connectivity MRI networks arise from subject motion. Neuroimage. 59: 2142–2154.

65. Smith SM, Nichols TE (2009): Threshold-free cluster enhancement: addressing problems of smoothing, threshold dependence and localisation in cluster inference. Neuroimage. 44: 83–98.

66. Winkler AM, Ridgway GR, Webster MA, Smith SM, Nichols TE (2014): Permutation inference for the general linear model. Neuroimage. 92: 381–397.

67. Esterman M, Tamber-Rosenau BJ, Chiu YC, Yantis S (2010): Avoiding non-independence in fMRI data analysis: leave one subject out. Neuroimage. 50: 572–576.

68. Osborne JW, Costello AB (2005): Best practices in exploratory factor analysis: four recommendations for getting the most from your analysis. Pan-Pacific Manag Rev. 2: 131–146.

69. Cadena EJ, White DM, Kraguljac N V., Reid MA, Lahti AC (2018): Evaluation of fronto-striatal networks during cognitive control in unmedicated patients with schizophrenia and the effect of antipsychotic medication. NPJ Schizophr. 4: 8.

70. Fornito A, Yung AR, Wood SJ, Phillips LJ, Nelson B, Cotton S, et al. (2008): Anatomic abnormalities of the anterior cingulate cortex before psychosis onset: an MRI study of ultra-high-risk individuals. Biol Psychiatry. 64: 758–765.

71. Fornito A, Yücel M, Wood SJ, Adamson C, Velakoulis D, Saling MM, et al. (2008): Surface-based morphometry of the anterior cingulate cortex in first episode schizophrenia. Hum Brain Mapp. 29: 478–489.

72. Bora E, Fornito A, Radua J, Walterfang M, Seal M, Wood SJ, et al. (2011): Neuroanatomical abnormalities in schizophrenia: a multimodal voxelwise meta-analysis and meta-regression analysis. Schizophr Res. 127: 46–57.

73. Fornito A, Yücel M, Patti J, Wood SJ, Pantelis C (2009): Mapping grey matter reductions in schizophrenia: an anatomical likelihood estimation analysis of voxel-based morphometry studies. Schizophr Res. 108: 104–113.

74. Reid MA, Stoeckel LE, White DM, Avsar KB, Bolding MS, Akella NS, et al. (2010): Assessments of function and biochemistry of the anterior cingulate cortex in schizophrenia. Biol Psychiatry. 68: 625–633.

75. Cassidy CM, Balsam PD, Weinstein JJ, Rosengard RJ, Slifstein M, Daw ND, et al. (2018): A perceptual inference mechanism for hallucinations linked to striatal dopamine. Curr Biol. 28: 503–514.e4.

76. Whitford V, O’Driscoll GA, Pack CC, Joober R, Malla A, Titone D (2013): Reading impairments in schizophrenia relate to individual differences in phonological processing and oculomotor control: evidence from a gaze-contingent moving window paradigm. J Exp Psychol Gen. 142: 57–75.

77. Butler PD, Zemon V, Schechter I, Saperstein AM, Hoptman MJ, Lim KO, et al. (2005): Early-stage visual processing and cortical amplification deficits in schizophrenia. Arch Gen Psychiatry. doi: 10.1001/archpsyc.62.5.495.

78. Papanastasiou E, Mouchlianitis E, Joyce DW, McGuire P, Banaschewski T, Bokde ALW, et al. (2018): Examination of the neural basis of psychoticlike experiences in adolescence during reward processing. JAMA Psychiatry. 1–9.

79. Lincoln TM, Dollfus S, Lyne J (2017): Current developments and challenges in the assessment of negative symptoms. Schizophr Res. 186: 8–18.

80. Carlsson A, Waters N, Carlsson ML (1999): Neurotransmitter interactions in schizophrenia--therapeutic implications. Biol Psychiatry. 46: 1388–1395.

81. Dazzan P, Morgan KD, Orr KG, Hutchinson G, Chitnis X, Suckling J, et al. (2004): The structural brain correlates of neurological soft signs in ÆSOP first-episode psychoses study. Brain. 127: 143–153.

82. Emsley R, Rabinowitz J, Torreman M, Schooler N, Kapala L, Davidson M, McGory P (2003): The factor structure for the Positive and Negative Syndrome Scale (PANSS) in recent-onset psychosis. Schizophr Res. 61: 47–57.

83. Walther S, Strik W (2012): Motor symptoms and schizophrenia. Neuropsychobiology. 66: 77–92.

84. Wang Y, Liu WH, Li Z, Wei XH, Jiang XQ, Geng FL, et al. (2016): Altered corticostriatal functional connectivity in individuals with high social anhedonia. Psychol Med. 46: 125–135.

85. Loranger AW (1984): Sex difference in age at onset of schizophrenia. Arch Gen Psychiatry. 41: 157–161.

86. Rössler W, Riecher-Rössler A, Angst J, Murray R, Gamma A, Eich D, et al. (2007): Psychotic experiences in the general population: a twenty-year prospective community study. Schizophr Res. 92: 1–14.

87. Satterthwaite TD, Wolf DH, Loughead J, Ruparel K, Elliott MA, Hakonarson H, et al. (2012): Impact of in-scanner head motion on multiple measures of functional connectivity: relevance for studies of neurodevelopment in youth. Neuroimage. 60: 623–632.

88. Ciric R, Wolf DH, Power JD, Roalf DR, Baum GL, Ruparel K, et al. (2017): Benchmarking of participant-level confound regression strategies for the control of motion artifact in studies of functional connectivity. Neuroimage. 154: 174–187.

